# Optogenetic dissection of actomyosin-dependent mechanics underlying tissue fluidity

**DOI:** 10.1101/2021.11.07.467642

**Authors:** R. Marisol Herrera-Perez, Christian Cupo, Cole Allan, Alicia B. Dagle, Karen E. Kasza

## Abstract

Rapid epithelial tissue flows are essential to building and shaping developing embryos. However, it is not well understood how the mechanical properties of tissues and the forces driving them to flow are jointly regulated to accommodate rapid tissue remodeling. To dissect the roles of actomyosin in the mechanics of epithelial tissue flows, here we use two optogenetic tools, optoGEF and optoGAP, to manipulate Rho/Rho-kinase signaling and actomyosin contractility in the germband epithelium, which flows via convergent extension movements during *Drosophila* body axis elongation. The ability to perturb actomyosin in the tissue allows us to analyze the effects of actomyosin on cell rearrangements, tissue tensions, and tissue mechanical properties. We find that either optogenetic activation or deactivation of Rho1 signaling and actomyosin contractility at the apical surface of the germband disrupts cell rearrangements and tissue-level flows. Rho1 activation leads to poorly oriented rearrangements that are associated with a redistribution of myosin II from the junctional to the medial-apical domain, whereas Rho1 deactivation leads to fewer, slower cell rearrangements that are associated with decreased junctional and medial myosin. By probing mechanical tensions in the tissue using laser ablation and inferring tissue mechanical properties from cell packings, we find that actomyosin influences *both* the anisotropic forces that drive tissue flow and the mechanical properties of the tissue that resist flow, leading to complex relationships between actomyosin activity and tissue fluidity. Moreover, our results link the subcellular distribution of myosin II to tissue tension and cell packings, revealing how junctional and medial myosin have differential roles in promoting and orienting cell rearrangements during tissue flow.

## INTRODUCTION

Within developing embryos, epithelial tissue sheets can dramatically deform to help lay out and shape the basic body plan of the animal. Many epithelial tissue movements are rapid and involve significant remodeling of the tissue through local cell rearrangements that alter the relative positions of cells within the tissue, similar to the deformation and flow of fluid materials (Herrera-Perez and Kasza, 2018; Tetley and Mao, 2018; Walck-Shannon and Hardin, 2014). Based on this similarity to fluid-like material deformation, the ability of a tissue to flow via cell rearrangements is often referred to as *tissue fluidity.* In physical models of epithelial tissues, cell rearrangements occur and the tissue accommodates flow if the energy barriers to rearrangement are small and/or the active driving forces are large (Bi et al., 2016, 2015, 2014; Duclut et al., 2021; Farhadifar et al., 2007; Popović et al., 2021; Staple et al., 2010). In contrast, if the energy barriers to rearrangement are large and the driving forces are small, cell rearrangements do not occur and the tissue resists flow in a solid-like manner. Thus, to understand the origins of tissue-scale flows, we must understand both the mechanical forces that drive flow and the mechanical properties of the tissue that determine how the tissue resists flowing in response to those forces. Recent studies suggest that the mechanical properties of tissues within developing embryos are regulated in space and time to allow (or prevent) rapid tissue remodeling and flow (Banavar et al., 2021; Jain et al., 2020; Mongera et al., 2018; Tetley et al., 2019; Wang et al., 2020). However, the molecular and cellular mechanisms involved in this regulation remain unclear.

Contractile networks of F-actin and non-muscle myosin II, jointly referred to as *actomyosin,* are a primary force-generating machinery in epithelial cells, with specific patterns of myosin localization and contractile activity correlating with tissue-level flow patterns (Heisenberg and Bellaïche, 2013; Herrera-Perez and Kasza, 2018; Kasza and Zallen, 2011; Martin, 2010). In addition, the mechanical behavior of the actomyosin cytoskeleton within cells has been implicated in controlling the mechanical properties of cells and the tissues in which they reside, with increased actomyosin-generated tension typically associated with stiffer cells and more solid-like tissues (Bi et al., 2015; Gardel et al., 2006; Wang et al., 2002; Zhou et al., 2009), which would be predicted to increase resistance to flow. Given these potential dual roles for actomyosin in regulating both tissue-scale driving forces and tissue mechanical properties, it is not well understood how patterns of actomyosin contractility might be regulated to tune tissue fluidity and overall tissue flow.

Body axis elongation in the *Drosophila* embryo is a powerful model system for studying the molecular and cellular mechanisms that contribute to epithelial tissue flows. During this morphogenetic event, the germband epithelium rapidly extends along the head-to-tail (anterior-posterior) body axis and narrows along the dorsal-ventral axis, a process driven by both cell rearrangements (Bertet et al., 2004; Blankenship et al., 2006; Farrell et al., 2017; Irvine and Wieschaus, 1994) and changes in cell shape (Butler et al., 2009; Collinet et al., 2015; Farrell et al., 2017; Lye et al., 2015). Planar polarized patterns of myosin II localization and activity are required for convergent extension of the germband tissue (Bertet et al., 2004; Blankenship et al., 2006; Fernandez-Gonzalez et al., 2009; Kasza et al., 2014; Munjal et al., 2015; Rauzi et al., 2010, 2008; Zallen and Wieschaus, 2004). The planar polarized myosin pattern generates anisotropic forces in the tissue that are thought to drive and orient cell rearrangement and directional tissue flow. Recent work from our group and others has demonstrated that the onset of rapid cell rearrangements in the germband is predicted by a solid-to-fluid transition in anisotropic vertex models of epithelial tissues (Wang et al., 2020), suggesting that the mechanical properties of the germband might be regulated to promote this rapid change in tissue shape. However, it is unclear what role actomyosin might play in regulating this change in germband tissue mechanical properties.

Here, we dissect the roles of actomyosin in tissue flows during *Drosophila* body axis elongation. This requires precise perturbation of actomyosin contractility patterns, specifically in the germband during body axis elongation, so that the mechanical behavior of neighboring regions of the embryo is not disturbed. Because this is not possible with traditional molecular genetics approaches, here we employed two previously described optogenetic tools, optoGEF and optoGAP, to manipulate the localization of upstream regulators of *Drosophila* Rho1 at the apical surface of the germband epithelium (Herrera-Perez et al., 2021). These tools are based on the CRY2/CIB1 heterodimerization system, which can be activated by blue-light illumination (Guglielmi et al., 2015; Idevall-Hagren et al., 2012; Izquierdo et al., 2018; Kennedy et al., 2010). The optoGEF tool is based on the full-length *Drosophila* RhoGEF2 protein and activates actomyosin contractility (Herrera-Perez et al., 2021). In contrast, optoGAP is based on the full-length *Drosophila* RhoGAP71E protein and deactivates actomyosin contractility (Herrera-Perez et al., 2021). Photoactivation via blue light illumination of the apical surface of the germband leads to rapid accumulation of the CRY2 component of the optogenetic tools at the apical cell membrane, allowing us to perturb actomyosin contractility rapidly at minute timescales and specifically in the germband tissue during body axis elongation (Herrera-Perez et al., 2021). Employing this optogenetics approach, we analyzed how perturbing actomyosin affects cell rearrangements (a measure of tissue fluidity) using live imaging and quantitative image analysis, how it affects mechanical forces in the tissue using laser ablation assays, and how it affects mechanical properties of the tissue by analyzing the structure of epithelial cell packings. Our findings indicate that actomyosin influences both the driving forces and tissue mechanical properties and reveal how distinct aspects of the actomyosin machinery promote cell rearrangement and tissue flow.

## RESULTS

### Optogenetic manipulation of Rho signaling disrupts cell rearrangements and tissue flow during convergent extension

To test how Rho1 signaling and actomyosin contractility influence tissue fluidity during convergent extension, we used light-gated optoGEF and optoGAP tools (Herrera-Perez et al., 2021) to manipulate Rho1/Rho-kinase signaling in the germband epithelium of the *Drosophila* embryo during body axis elongation (Fig 1 *A,B*) and analyzed the effects on cell rearrangements and tissue flow. We generated embryos expressing the optoGEF or optoGAP tool using the Gal4/UAS system and illuminated the apical surface of the ventrolateral region of the germband epithelium with blue light (488 nm), beginning at the onset of body axis elongation, which we define as *t* = 0. We captured time-lapse confocal movies and used quantitative image analysis to quantify the effects on the number of cell rearrangements and overall tissue elongation.

**Figure 1.**
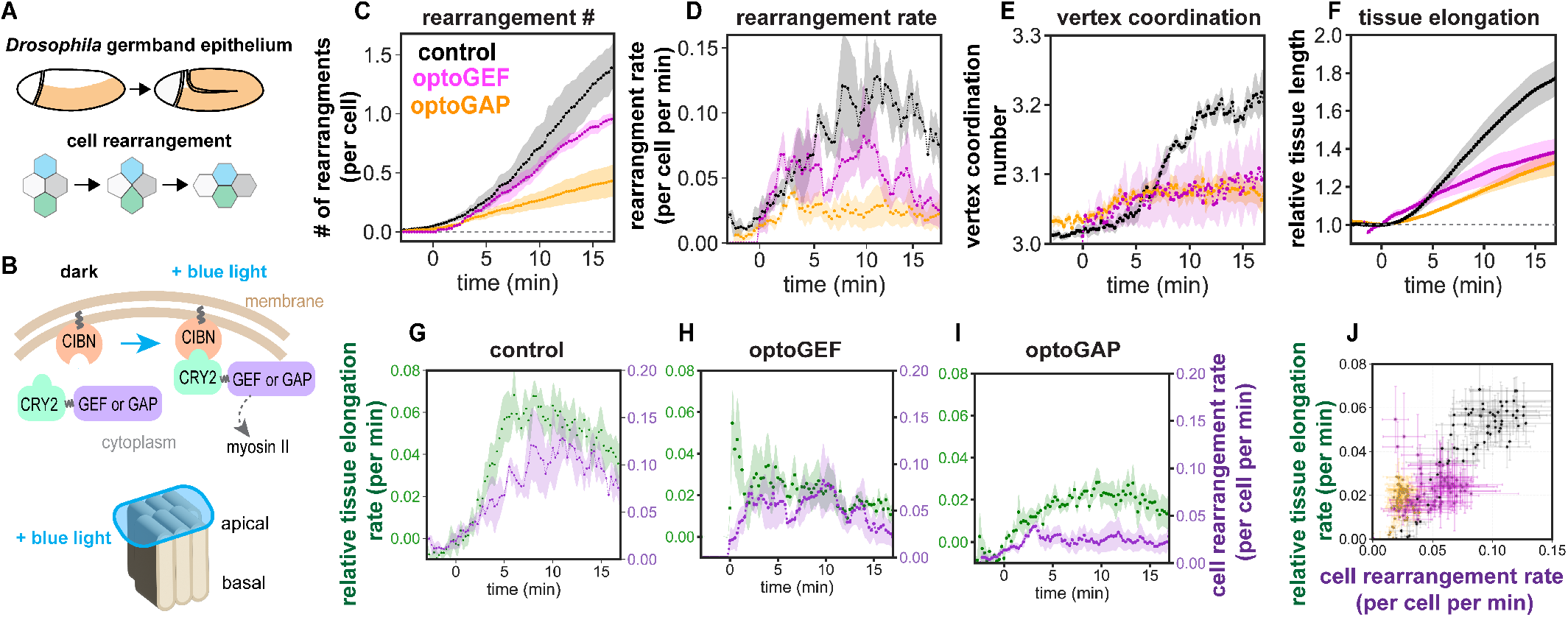
Optogenetic manipulation of Rho signaling tunes cell rearrangements rates and tissue flow during *Drosophila* body axis elongation. **(A)** *Top*, Schematic of *Drosophila* body axis elongation. The germband epithelial tissue (orange) narrows and elongates the embryonic body axis in a convergent extension movement driven by oriented cell rearrangements. *Bottom,* Schematic of an oriented cell rearrangement. **(B)** *Top,* Schematic of optogenetic tools. CIBN is targeted to the cell membrane. The light-sensitive PHR domain of CRY2 is fused to upstream myosin II regulators of the Rho/Rho-kinase pathway, RhoGEF2 (GEF) or RhoGAP71E (GAP). Under blue light illumination, CRY2 dimerizes with CIBN and the regulators accumulate at the cell membrane. *Bottom*, Blue-light illumination of the apical surface of the epithelium. **(C)** Number of cell rearrangements initiated in control, optoGEF, and optoGAP embryos, including both T1 processes and higher-order rosette rearrangement events. **(D)** Instantaneous rate of rearrangements per cell per minute. **(E)** Cell vertex coordination number represents the average number of cells meeting at a common vertex in the tissue. **(F)** Elongation of the germband tissue during convergent extension quantified by PIV analysis of time-lapse confocal movies of embryos. The length of the tissue was normalized to the value at *t* = 0. **(G-I)** Instantaneous rates of tissue elongation and cell rearrangements over time for control (G), optoGEF (H), and optoGAP (I) embryos. **(J)** Correlation between the instantaneous rates of tissue elongation and cell rearrangement. Each data point represents the mean between embryos at an individual time point. The mean ± SEM between embryos is shown, n=3-5 embryos per genotype.

First, we analyzed the effects of optogenetic perturbation of Rho1 signaling on the number of cell rearrangements that occur in the tissue as a metric for *tissue fluidity,* the ability of a tissue to remodel and flow. In control embryos expressing only the CIBN-pmGFP component of the optogenetic system, there were 1.25 ± 0.19 cell rearrangements initiated per cell in the germband during the first 15 min of axis elongation, similar to that seen in wild-type embryos (Farrell et al., 2017; Kasza et al., 2019, 2014; Paré et al., 2014; Simões et al., 2010) (Fig 1*C*). In contrast, under blue-light illumination of the ventrolateral region of the germband, there were 0.93 ± 0.05 rearrangements per cell in optoGEF embryos (*P* = 0.04 compared to controls), and 0.40 ± 0.12 rearrangements per cell in optoGAP embryos (*P* = 0.03 compared to controls) (Fig 1*C*). Consistent with reduced cell rearrangements, which involve transient cell configurations in which four or more cells meet at a common vertex, photoactivated optoGEF and optoGAP embryos displayed a decrease in average vertex coordination number in the germband (3.07± 0.04, *P* = 0.05 for optoGEF and 3.07± 0.02, *P* = 0.04 for optoGAP at *t* = 15 min) compared to control embryos (3.18 ± 0.01 at *t* = 15 min) (Fig. 1*E*). Taken together, these results show that cell rearrangements are reduced when Rho1 activity is *either* increased (optoGEF) or decreased (optoGAP) in the germband, although the defects are more severe when Rho1 activity is decreased. This is consistent with key roles for Rho1 signaling and actomyosin contractility in controlling tissue fluidity.

Next, we analyzed how these changes in tissue fluidity influence macroscopic tissue flow. In control embryos expressing only the CIBN-pmGFP component of the optogenetic system, the germband tissue elongated 1.71 ± 0.09-fold along the head-to-tail axis (Fig. 1*F*, *t* = 15 min), similar to in wild-type embryos (Farrell et al., 2017; Kasza et al., 2014; Simões et al., 2010). In contrast, the germband in optoGEF embryos elongated 1.37 ± 0.09-fold (*P* = 0.06), while the germband in optoGAP embryos elongated 1.29 ± 0.05-fold (*P* = 0.02) (Fig. 1*F*), consistent with the notion that cell rearrangements mediate tissue flow during convergent extension. Interestingly, tissue-level elongation was similar in both the optoGEF and optoGAP perturbations, despite there being more rearrangements initiated in optoGEF compared to optoGAP embryos, suggesting that other factors might be differentially altered by these perturbations, such as cell shape or the spatial orientation of cell rearrangements.

To take a closer look at cell rearrangements and tissue flow, and the relationships between them, we analyzed the instantaneous rates of cell rearrangement and tissue elongation. In control embryos, the rate of relative tissue elongation increases during the first 10 min of axis elongation to a value of 0.05 ± 0.01 min^-1^ at *t* = 10 min, followed by a decrease during the rest of the process (Fig. 1*G*). The cell rearrangement rate follows a similar trend, reaching a value of 0.12 ± 0.04 cell^-1^ min^-1^ at *t* = 10 min, but with a delay relative to tissue elongation (Fig. 1*G*). This delay is possibly associated with cell shape changes during this time that can contribute to tissue-level elongation (Butler et al., 2009; Farrell et al., 2017; Lye et al., 2015). In optoGEF and optoGAP embryos, the rates of relative tissue elongation and cell rearrangement were both decreased (Fig. 1*H,I*). At *t* = 10 min, the rate of tissue elongation was reduced to 0.02 ± 0.01 min^-1^ in optoGEF (*P* = 0.09) and 0.02 ± 0.06 min^-1^ in optoGAP embryos (*P* = 0.03). At *t* = 10 min, the rate of cell rearrangements was 0.08 ± 0.03 cell^-1^ min^-1^ in optoGEF (*P* = 0.45) and 0.04 ± 0.02 cell^-1^ min^-1^ in optoGAP embryos (*P* = 0.09). The instantaneous rates of tissue elongation and cell rearrangement are correlated in all embryo groups (Fig. 1*J*, correlation coefficient for control=0.93, optoGEF=0.58, optoGAP=0.71), consistent with a central role for tissue fluidity in promoting tissue elongation. Notably, however, the rate of cell rearrangement was the least predictive of tissue elongation rate for the optoGEF perturbation.

Taken together, these results demonstrate that optogenetic perturbation of Rho1 signaling and actomyosin activity at the apical surface of the germband epithelium disrupts cell rearrangements, decreasing the ability of the tissue to remodel and flow like a fluid during convergent extension.

### Rho signaling influences predicted tissue mechanical properties, which characterize resistance to tissue flow

To understand why a tissue accommodates (or resists) rapid remodeling and flow, one must consider both the mechanical forces that act on the tissue to drive flow and the tissue mechanical properties that capture how the tissue resists flowing under those forces. Planar polarized patterns of actomyosin in the germband generate contractile forces that are thought to be driving forces for tissue flow (Bertet et al., 2004; Blankenship et al., 2006; Fernandez-Gonzalez et al., 2009; Kasza et al., 2014; Munjal et al., 2015; Rauzi et al., 2008; Zallen and Wieschaus, 2004). In addition, actomyosin contractility has also been proposed to influence the mechanical properties of epithelial tissues, with increased actomyosin-generated tension predicted to increase the resistance of the tissue to flow (Bi et al., 2015). Because actomyosin potentially plays dual roles in influencing how tissues flow, we wondered if effects on driving forces and/or tissue mechanical properties could explain the observed reductions in tissue fluidity when Rho activity is perturbed.

First, we tested if tissue mechanical properties are altered when the elongating germband is photoactivated in optoGEF or optoGAP embryos. Because direct mechanical measurements to probe the mechanical response of the germband remain a significant experimental challenge, we gained insight from predictions of anisotropic vertex models of epithelial tissues. Recent work from our group and others has demonstrated that the fluid-like versus solid-like behavior of epithelia can be predicted from analysis of cell packings in the tissue, which can be extracted from images of cell outlines (Jain et al., 2020; Mitchel et al., 2020; Park et al., 2015; Wang et al., 2020). In particular, for anisotropic tissues like the germband, two metrics of cell packings in a tissue, the average cell shape index 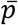 and cell shape alignment index *Q*, are sufficient to classify a given packing as either solid-like or fluid-like (Wang et al., 2020). This link between cell shapes and tissue mechanics can be understood from theoretical vertex models of epithelia in which cell packings determine the energy barriers to cell rearrangement, thereby controlling whether the model tissue behaves in a solid-like (high energy barriers, few rearrangements) or fluid-like (low energy barriers, many rearrangements) manner. To gain insight into how Rho signaling influences germband tissue mechanical properties, we analyzed cell shape and alignment in photoactivated control, optoGEF, and optoGAP embryos.

We quantified how the average cell shape index, 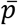 (cell perimeter divided by the square root of cell area, averaged across the cells in the tissue), and the cell shape alignment index, *Q* (average cell elongation across the tissue, similar to a nematic order parameter but modulated by the extent of cell elongation), varied over time following blue-light illumination of the ventrolateral region of the germband beginning at the onset of axis elongation (Fig. 2*A-C*). In control embryos expressing only the CIBN-pmGFP component of the optogenetic system, the average cell shape index, 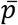, increased over time and the cell shape alignment index, *Q*, showed a peak around *t* = 2 min (Fig. 2*A-C*), similar to in wild-type embryos (Wang et al., 2020). Comparing these results to the predictions of the anisotropic vertex model (Wang et al., 2020) suggests that the germband tissue in control embryos starts out near the predicted solid-fluid transition line in *p-Q* parameter space and over time moves deeper into the predicted fluid-like region during tissue elongation (Fig. 2*D,E*), consistent with tissue properties becoming more fluid-like to minimize resistance to flow during axis elongation.

**Figure 2.**
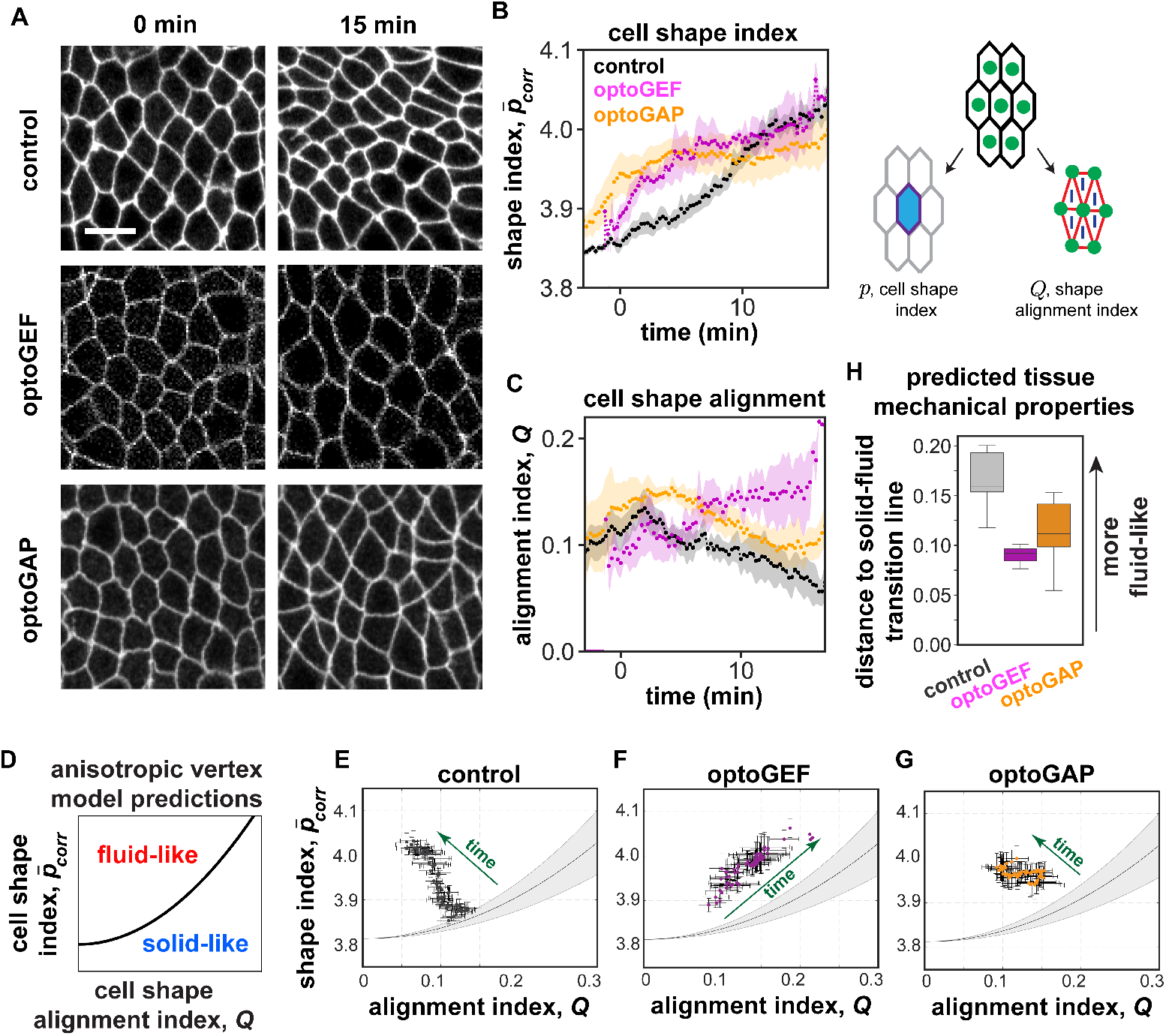
Cell packings and tissue mechanics in optogenetically manipulated germband tissue. **(A)** Stills from confocal movies showing cell packings during axis elongation. Anterior, left. Ventral, down. Bar, 10 μm. **(B)** The corrected average cell shape index 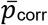 during convergent extension in photoactivated control, optoGEF, and optoGAP embryos. 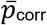 is calculated as described in the Materials and Methods section. The mean ± SEM between embryos is shown, n=3-5 embryos per genotype. **(C)** The cell shape alignment index *Q* during convergent extension in photoactivated control, optoGEF, and optoGAP embryos. *Q* is calculated as described in the Materials and Methods section. The mean ± SEM between embryos is shown, n=3-5 embryos per genotype. **(D)** The corrected cell shape index and cell shape alignment can be used to predict tissue mechanical behavior using an anisotropic vertex model (Wang et al., 2020). **(E-G)** Relationship between the corrected averaged cell shape index and cell shape alignment in the germband for control (E), optoGEF (F), and optoGAP (G) embryos. Each data point represents the mean between embryos at an individual time point, n=3-5 embryos per genotype. Solid line represents the predicted solid-fluid transition line, shaded area represents the range of predicted values for the transition from prior vertex model simulations (see Materials and Methods section). **(H)** Distance to the predicted solid-fluid transition line, evaluated during the rapid phase of tissue elongation (*t* = 15 min).

In contrast, cell packings in the germband of photoactivated optoGEF and optoGAP embryos were altered in distinct ways, leading to differences in predicted tissue mechanical properties. In photoactivated optoGAP embryos, the average cell shape index 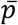 was increased compared to controls during the first 10 min of the process, and the alignment index *Q* showed similar trends as controls with somewhat elevated values of alignment compared to controls throughout the process (Fig. 2*A-C*). Comparing these results to the vertex model predictions suggests that in optoGAP embryos, the germband starts out near the predicted solid-fluid transition line, similar to controls, but does not move as deeply into the predicted fluid-like region (Fig. 2*D,G*), possibly consistent with less fluid-like behavior and more resistance to flow.

In photoactivated optoGEF embryos, although the average cell shape index 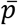 displays similar trends as in the optoGAP case, the alignment index *Q* shows qualitatively different behavior compared to control or optoGAP embryos (Fig. 2*A-C*). Indeed, the cell shape alignment *Q* remained high throughout the process, and did not relax back down to low levels as observed in controls. Comparing these results to the vertex model predictions suggests that the germband in optoGEF embryos again starts out near the predicted solid-fluid transition line, similar to controls, but takes a path through the *p-Q* parameter space over time that follows along the transition line instead of relaxing deeper into the fluid-like region (Fig. 2*D,F*), possibly consistent with significantly less fluid-like behavior and more resistance to flow.

To compare the predicted tissue mechanical properties in the control, optoGEF, and optoGAP groups, we measured the distance to the predicted solid-fluid transition line in the *p-Q* parameter space. During the rapid phase of tissue elongation, the distance to the transition line was reduced in optoGAP (*P =* 0.05) and significantly reduced in optoGEF (*P =* 0.005) compared to in control embryos (Fig. 2*D,H*). Taken together, these findings suggest that tissue mechanical properties, which capture how a tissue resists flowing under an applied force, are altered by either activation (optoGEF) or inactivation (optoGAP) of Rho signaling in the germband. In both cases, the germband is predicted to be less fluid-like than in control embryos, which is consistent with the observed reduction in cell rearrangements and tissue flow. However, these results are not sufficient to explain why we observe fewer cell rearrangements in optoGAP compared to optoGEF embryos, despite the prediction of even less fluid-like behavior in the optoGEF case.

### Rho signaling influences the planar polarized patterns of myosin localization at cell junctions

We wondered if differences in the mechanical forces driving tissue flow might help to account for the cell rearrangement rates and tissue flows in optoGEF and optoGAP embryos. During germband extension, Rho-kinase (Rok) is necessary for the formation of planar polarized patterns of myosin II localization and activity (Bertet et al., 2004; Kasza et al., 2014; Kerridge et al., 2016; Simões et al., 2014, 2010) that are thought to provide driving forces for cell rearrangements and tissue flow during axis elongation (Bertet et al., 2004; Blankenship et al., 2006; Fernandez-Gonzalez et al., 2009; Kerridge et al., 2016; Rauzi et al., 2010, 2008; Zallen and Wieschaus, 2004). To test if myosin-generated driving forces are disrupted, we examined Rok and myosin II localization in photoactivated optoGEF and optoGAP embryos (Fig. 3*A-F*).

**Figure 3.**
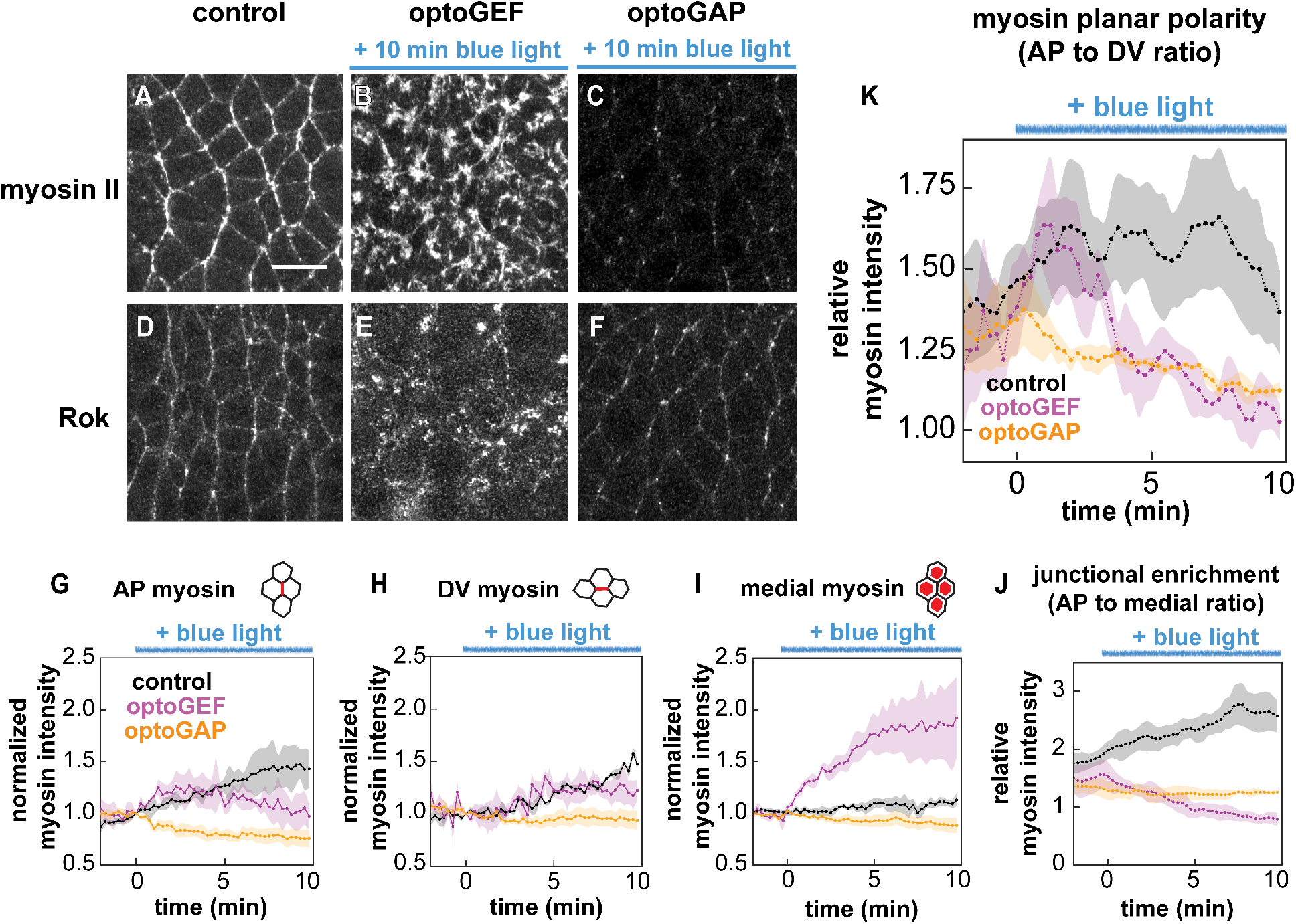
Myosin II and Rok localization patterns in optogenetically manipulated germband tissue. **(A-F)** Stills from movies of epithelial germband cells during *Drosophila* body axis elongation showing myosin II and Rho-kinase (Rok) localization in control, optoGEF, and optoGAP embryos after continuous activation of the tool for 10 min. Bar, 10 μm. Maximum intensity projection of z-slices in the most apical 5 μm. Myosin II (myosin) was visualized using an mCherry-tagged myosin regulatory light chain (*sqh*) transgene. Rok was visualized using a GFP-tagged Rho-kinase^K116A^ transgene under the myosin regulatory light chain (*sqh*) promoter. **(G-H)** Mean junctional myosin intensity before and during tool activation beginning at *t* = 0 (duration of blue light exposure indicated by blue line) at AP (*G*) and DV (*H*) junctions. Both optoGEF and optoGAP embryos exhibit decreased junctional myosin accumulation compared to controls, n=3-4 embryos, > 30 junctions/embryo. **(I)** Mean myosin intensity in the medial-apical domain of cells over time. The optoGEF embryos exhibit medial myosin accumulation, while optoGAP embryos show decreased medial myosin following blue light illumination beginning at *t* = 0. n=3-4 embryos, >10 cells/embryo. **(J)** Ratio of mean myosin intensities at AP junctions compared to in the medial-apical domain. Ratios for each embryo at each time point were calculated, and the mean ± SEM between embryos is shown. **(K)** Ratio of mean myosin intensities at AP compared to at DV junctions. Myosin planar polarity was disrupted in optoGEF and optoGAP embryos. Ratios for each embryo at each time point were calculated, and the mean ± SEM between embryo is shown.

First, we assessed the effects of optogenetic manipulation of Rho1 signaling on Rok localization using a GFP-tagged Rho-kinase^K116A^ transgene (Simões et al., 2014). In photoactivated control embryos, Rok localized at cell-cell contacts with enhanced accumulation at “vertical” edges between anterior and posterior cell neighbors (AP edges) compared to “horizontal” edges between dorsal and ventral cell neighbors (DV edges) (Fig. 3*D*), similar to in wild-type embryos (Simões et al., 2014, 2010). In photoactivated optoGEF embryos, the localization pattern of Rok was strongly disrupted, with Rok accumulating in clusters in the medial-apical domain of cells (Fig. 3*E*). In contrast, in photoactivated optoGAP embryos, Rok localized at cell-cell contacts, similar to in controls, but the overall levels were decreased and the planar polarized pattern appeared to be disrupted (Fig. 3*F*). Thus, perturbations to Rho1 in optoGEF and optoGAP embryos altered the localization pattern of Rok in the germband.

Next, to analyze how optogenetic recruitment of Rho1 regulators at the apical surface of the germband influences myosin II localization patterns, we performed time-lapse confocal imaging of myosin II in optoGEF and optoGAP embryos. We visualized myosin II using an mCherry-tagged myosin regulatory light chain. In optoGEF embryos, we observed that following blue light exposure starting at *t* = 0 to recruit RhoGEF2 to the cell membrane, myosin rapidly accumulated at the medial-apical cortex of cells (Fig. 3*B*), with a 22 ± 4% increase in medial myosin within 1 min and a 66 ± 14% increase over 5 min (Fig. 3*I*), compared to 3 ± 2% in 1 min and 9% ± 2% in 5 min in control embryos (Fig. 3*A,I*). This increase in medial myosin is consistent with the observed changes in Rok localization (Fig. 3*E*). The rapid increase in medial myosin in optoGEF embryos was accompanied by more gradual changes in junctional myosin levels. Myosin localization at AP and DV cell junctions initially increased, following the same trends as in control embryos, before beginning to decrease 3-5 min after the start of blue light exposure (Figs. 3*G-H*), which was delayed compared to the nearly immediate medial accumulation of myosin (Fig. 3*I*). Together, these changes in myosin localization produced a strong decrease in myosin planar polarity after 3 min of blue light exposure (Fig. 3*K*) and led to the formation of a radial pattern of myosin reminiscent of those found in tissues exhibiting apical constriction and invagination.

In contrast, photoactivation of optoGAP in the germband epithelium resulted in an immediate but gradual decrease in both medial and junctional myosin (Fig. 3*C,G-I*), demonstrating that ectopic RhoGAP71E cell membrane recruitment is sufficient to decrease cortical localization of myosin, which is consistent with the observed changes in Rok localization (Fig. 3*F*) and the role of RhoGAPs in promoting the inactive state of Rho1. The majority of the reduction in cortical myosin occurred over a timescale of ~5 min and was associated with an almost immediate decrease in myosin planar polarity compared to in control embryos (Fig. 3*K*). Thus, myosin planar polarity was disrupted in both optoGEF and optoGAP perturbations.

### Rho signaling influences the anisotropic forces at cell junctions thought to drive tissue flow

Next, we tested if the altered myosin localization patterns in photoactivated optoGEF and optoGAP embryos resulted in changes to the anisotropic mechanical forces in the tissue, which are thought to be a driving force for germband tissue flow. We used a laser to ablate individual junctional domains at the contacts between neighboring cells (Fig. 4*A,B*) (Farhadifar et al., 2007; Fernandez-Gonzalez et al., 2009; Herrera-Perez and Kasza, 2019; Hutson et al., 2003; Rauzi et al., 2008). Retraction of the tissue away from the ablation site reflects the mechanical tension in the tissue prior to ablation, with the initial retraction velocity being proportional to tension and inversely related to viscous drag. The decreased myosin planar polarity in photoactivated optoGEF and optoGAP embryos would be predicted to result in reduced anisotropy in forces along AP compared to DV cell junctions.

**Figure 4.**
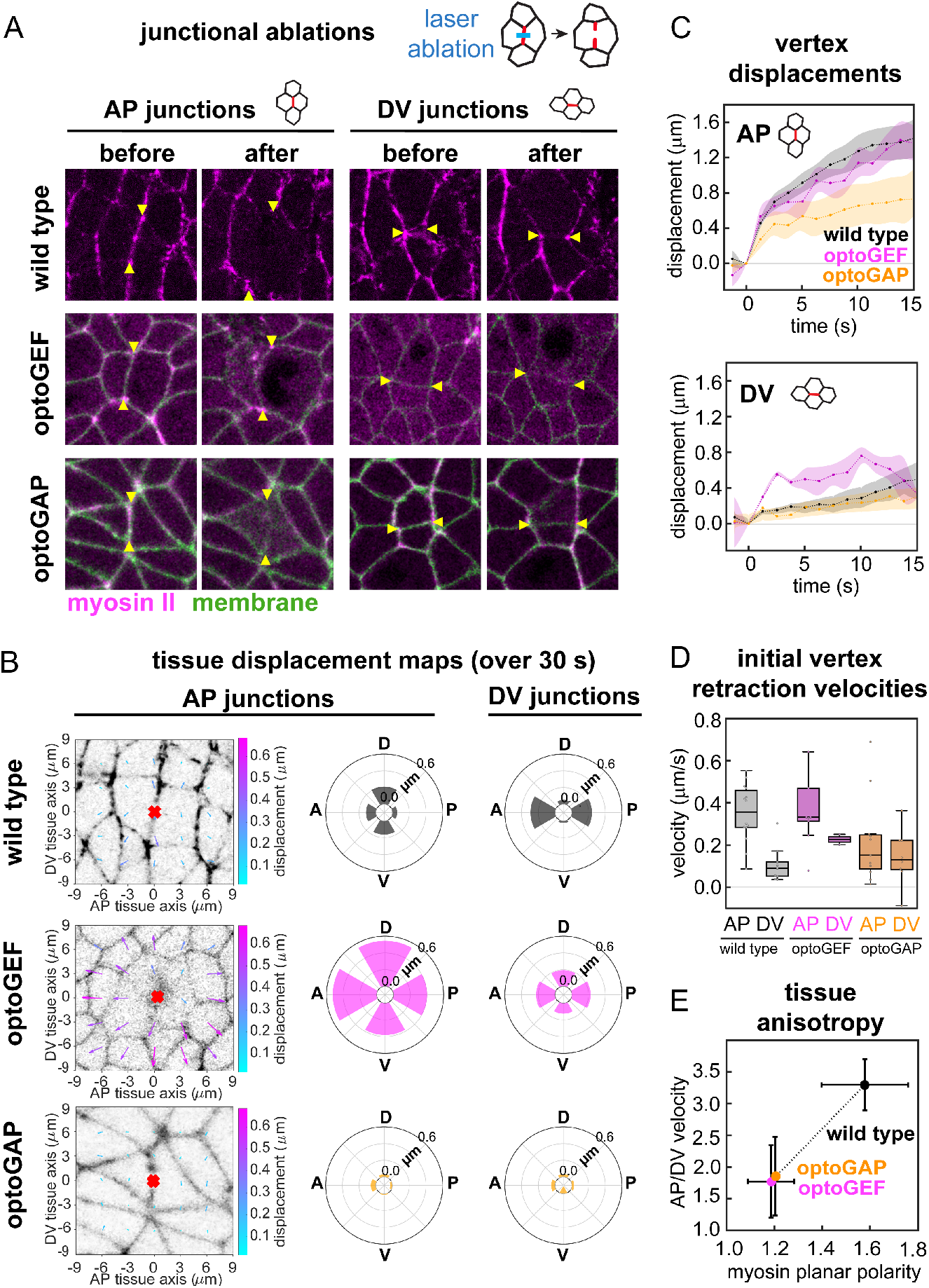
Ablations at cell-cell junctions in optogenetically manipulated germband tissue. **(A)** Cells before (left) and 1 min after (right) ablation. Myosin II (magenta), CIBN-pmGFP (green). Arrowheads indicate the vertices connected to the junction in which the cut was made. Anterior, left. Ventral, down. Image size, 20 x 20 μm. **(B)** Maps of displacements in the surrounding tissue in the 30 s following ablations. The tissue in a 174 μm^2^ region surrounding the ablation point (red mark) was analyzed by PIV analysis following ablation. Radial plots show the mean total displacement, averaged over the regions in 60° angular bins, along the anterior, posterior, dorsal, or ventral tissue axes, n = 3-1 1 ablations per condition. **(C)** Vertex displacements following ablations at AP or DV junctions. Mean ± SEM between junctions is shown, n=10-15 AP, 3-10 DV junctions per condition. **(D)** Initial vertex retraction velocities after ablation. **(E)** Mechanical anisotropy was reduced in optoGEF and optoGAP embryos and correlates with myosin planar polarity at 5 min after optogenetic activation (c.f. Fig. 3*K*). Mechanical anisotropy was calculated as the ratio of mean AP to mean DV peak retraction velocities, n= 3-15 edges per condition.

In wild-type control embryos, the mean initial retraction velocities were 0.36 ± 0.04 μm/s at AP junctions and 0.11 ± 0.03 μm/s at DV junctions (Fig. 4*C,D*), producing a significant mechanical anisotropy in the tissue (comparison between AP and DV velocities, *P* = 9 ×10^-6^, *t-*test) (Fig. 4*E*), consistent with previous studies of mechanical anisotropy in the germband of wild-type embryos (Fernandez-Gonzalez et al., 2009; Kasza et al., 2014; Rauzi et al., 2008). In contrast, the mechanical anisotropies in photoactivated optoGEF and optoGAP embryos were reduced (Fig. 4*C-E*), such that the differences between velocities at AP junctions and at DV junctions were less significant (*P* = 0.04, *P* = 0.16 respectively, *t-*test), consistent with the reduced myosin planar polarity in these tissues.

Taken together, these results demonstrate that optogenetic activation or inactivation of Rho signaling nearly abolishes myosin planar polarity across the germband and disrupts the anisotropic contractile tensions associated with myosin activity at cell junctions, helping to explain why cell rearrangements are reduced in both perturbations. However, these results do not fully explain the differences in the defects observed in the optoGEF and optoGAP embryos. In optoGAP embryos, the germband is predicted to be less fluid-like than in wild-type embryos, and the near abolishment of planar polarized forces from myosin activity results in fewer cell rearrangements and reduced tissue flow. However, in optoGEF embryos, myosin planar polarity is also nearly abolished, and the tissue is predicted to be even less fluid-like, but we observe significantly more cell rearrangements initiated than in the optoGAP case. Furthermore, the cell rearrangements in optoGEF embryos appear to be less productive at contributing to convergent extension, as the final extent of germband tissue elongation is similar in optoGEF and optoGAP embryos.

### Rho signaling tunes myosin-dependent forces arising in the medial-apical domain of cells during convergent extension

We wondered what additional features of cell behaviors might contribute to these distinct defects in the germband of photoactivated optoGEF and optoGAP embryos. One notable feature of myosin II localization in the optoGEF embryos is the significantly enhanced accumulation of myosin at the medial-apical domain of cells (c.f. Fig 3*B,I*). To test how this enhanced medial-apical myosin in optoGEF embryos influences mechanical tensions in the tissue that might contribute to cell rearrangements, we used a laser to ablate myosin in the medial-apical domains of individual cells (Figs. 5*A,B*).

**Figure 5.**
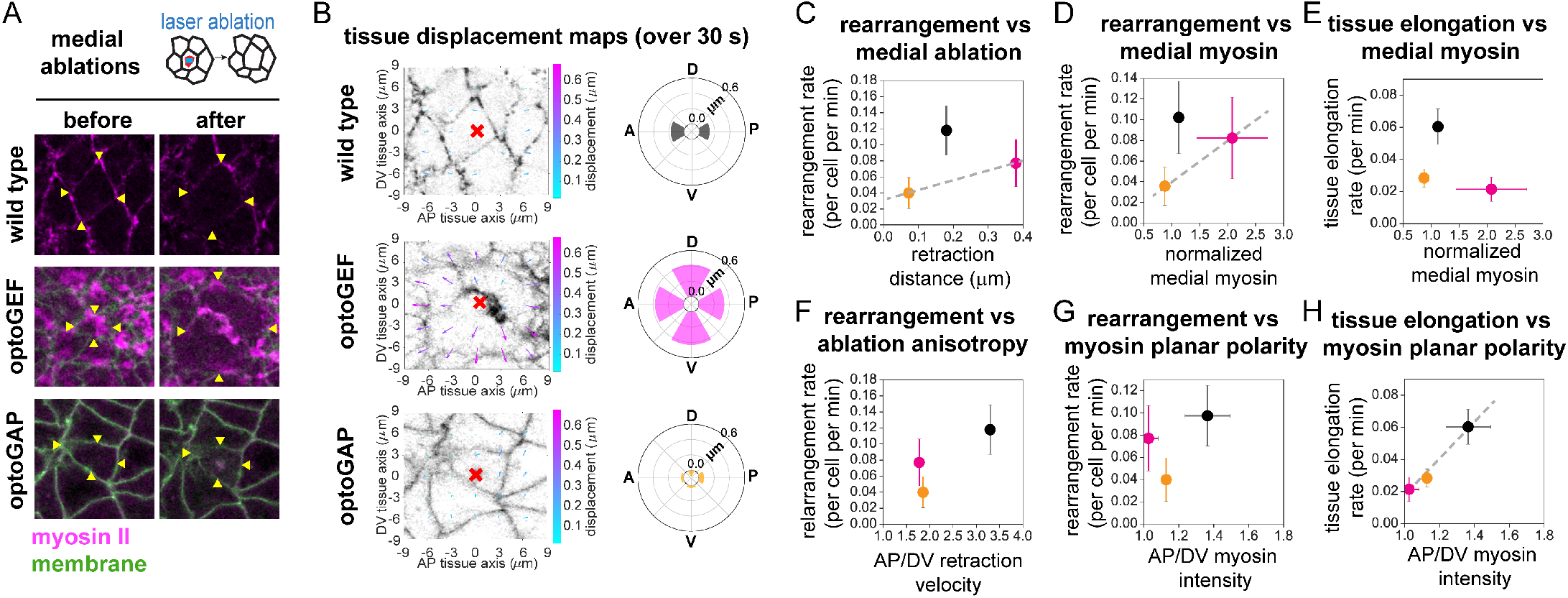
Ablations in the medial-apical domain of cells in optogenetically manipulated germband tissue. **(A)** Cells before (left) and 1 min after (right) ablation. Myosin II (magenta), CIBN-pmGFP (green). Arrowheads indicate the vertices connected to the cell in which the cut was made. Anterior, left. Ventral, down. Image size, 20 μm. **(B)** Maps of displacements in the surrounding tissue in the 30 s following ablations. The tissue in a 174 μm^2^ region surrounding the ablation point (red mark) was analyzed by PIV analysis following ablation. Radial plots show the mean total displacement, averaged over the regions in 60° angular bins, along the anterior, posterior, dorsal, or ventral tissue axes, n = 3-11 ablations per condition. **(C-H)** Relationships among cell rearrangement rates, tissue elongation rates, myosin patterns, and ablation results during convergent extension in photoactivated control (*black*), optoGEF (*magenta*), and optoGAP (*orange*) embryos (10 min time point). **(C-E)** Relationships between medial-apical myosin properties and tissue behaviors: (C) rearrangement rate vs. retraction distance at 30 s after ablation of medial-apical myosin, (D) rearrangement rate vs. normalized medial-apical myosin intensity, (E) tissue elongation rate vs. normalized medial-apical myosin intensity. Dashed lines, guides to the eye. **(F-H)** Relationships between planar polarized junctional myosin properties and tissue behaviors: (F) rearrangement rate vs. AP to DV ratio of peak retraction velocities after ablation of junctional myosin, (G) rearrangement rate vs. ratio of myosin intensity at AP and DV cell edges, (H) tissue elongation rate vs. ratio of myosin intensity at AP and DV cell edges.

In optoGEF embryos after 5 min of photoactivation, ablation in the medial-apical domain in a single cell led to dramatic retraction of the surrounding tissue along both the AP and DV axes (Fig. 5*A,B*), which was significantly enhanced compared to medial ablations in wild-type embryos (*P* = 0.018) and optoGAP embryos (*P* = 0.009), in which there was less medial myosin present (Fig. 5*A*). In addition, tissue retraction in optoGEF embryos was more isotropic than in controls. In contrast, medial ablations in photoactivated optoGAP embryos, resulted in retractions more similar in magnitude to control embryos (*P* = 0.39) (Fig. 5*A,B*). These results indicate that the myosin accumulating at the apical cortex of germband cells in optoGEF embryos is actively generating contractile forces contributing to mechanical tension in the tissue and that these tensions are more isotropic than in control embryos.

In a prior study, our group demonstrated that the medial-apical myosin in activated optoGEF cells shows dynamic pulsing behavior that is correlated with enhanced apical cell area fluctuations (Herrera-Perez et al., 2021). Here, we find that in the absence of strongly planar polarized junctional myosin in optoGEF and optoGAP embryos, cell rearrangement rates increase with tissue tension associated with medial myosin (Fig. 5*C*) and with medial myosin levels (Fig. 5*D*), even though strong junctional myosin planar polarity patterns are essential for harnessing cell rearrangement for tissue-level shape changes (Fig. 5*E-H*). Taken together, our findings suggest that Rho signaling tunes pulsatile contractile forces originating from the medial-apical domain of cells in the germband and that these forces promote cell rearrangement. This is consistent with recent studies highlighting a potential role for active tension fluctuations in promoting cell rearrangement by helping cells overcome the collective energy barrier associated with cell shape changes during this process (Duclut et al., 2021; Kim et al., 2021; Krajnc, 2020). Such a mechanism is consistent with the increased cell rearrangement rates in the optoGEF compared to optoGAP perturbations.

### Optogenetic manipulation of Rho disrupts distinct aspects of cell rearrangements

To gain further insight into how these perturbations to tissue mechanical properties and myosin-dependent driving forces influence cell rearrangements and tissue fluidity, we analyzed three distinct steps in cell rearrangement: cell edge contraction, vertex resolution, and new cell edge growth. We hypothesized that manipulation of Rho1, by increasing Rho1 activity (optoGEF) or decreasing Rho1 activity (optoGAP), might affect the rate or spatial organization of these steps in cell rearrangement, leading to the observed changes in cell rearrangement rates and tissue-level flow.

First, we examined whether activation of optoGEF or optoGAP across the tissue altered the speed of cell rearrangements that were initiated. In photoactivated optoGAP embryos, the median speed of contraction of AP or “vertical” cell edges between anterior and posterior cell neighbors (1.12 μm/min) was reduced compared to in control embryos (1.48 μm/min) (*P* = 0.013) (Fig. 6*A*), and the median rate for a high order vertex to resolve to form a DV cell edge was also reduced (0.11 min^-1^) compared to in control embryos (0.35 min^-1^) (*P* =0.02) (Fig. 6*B,F,H*). In addition, the median rate of growth of newly formed DV cell edges was decreased in optoGAP (0.93 μm/min) compared to in control embryos (1.26 μm/min) (*P*=0.03) (Fig. 6*C*), consistent with a role for actomyosin contractility both in edge contraction and in vertex resolution (Bertet et al., 2004; Blankenship et al., 2006; Collinet et al., 2015; Fernandez-Gonzalez et al., 2009; Kasza et al., 2019, 2014; Rauzi et al., 2008; Yu and Fernandez-Gonzalez, 2016). In contrast, the speeds of cell rearrangements were not significantly faster in photoactivated optoGEF embryos compared to in controls (Fig. 6*A-C,G*), suggesting that the increased myosin activity at the cell cortex in optoGEF embryos was not translated into significantly faster cell rearrangements once initiated.

**Figure 6.**
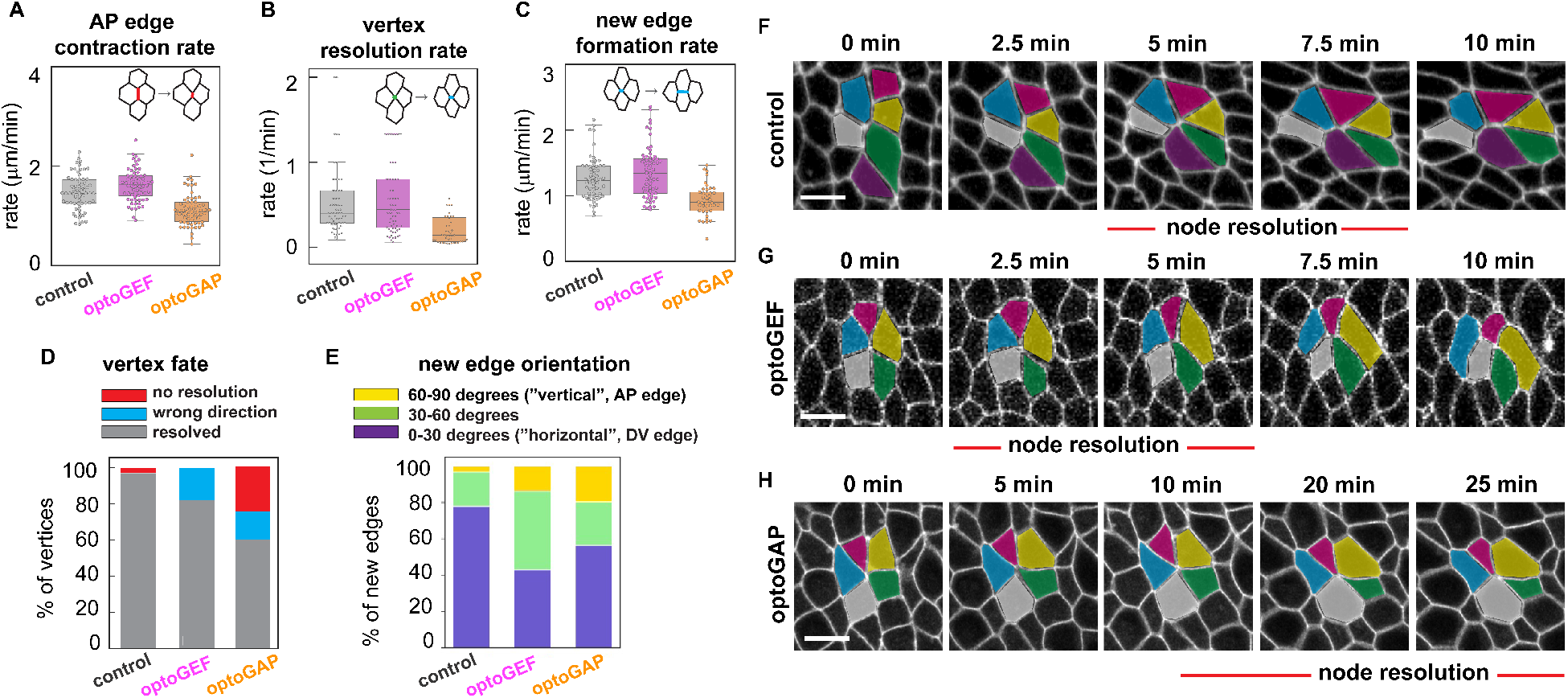
Cell rearrangement speed and spatial organization in optogenetically manipulated germband tissue. **(A-C)** AP cell edge contraction rate, vertex resolution rate, and new edge formation rate are reduced in optoGAP embryos compared to control embryos. Each data point represents a cell rearrangement initiated by contraction of an AP edge. n=10-25 edges/embryo from 3-5 embryos (n=48-69 edges per genotype). See details in the Materials and Methods section. **(D)** The fate of vertices formed through contraction of AP cell edges. The optoGAP and optoGEF embryos showed an increased percentage of vertices that did not resolve or aberrantly resolved to form an AP edge. **(E)** The orientation of newly formed cell edges. While the majority of the new edges in control embryos correspond to edges oriented parallel to the anterior-posterior axis of the embryo (orientation 0-30°), optoGEF embryos showed an increase in edges with diagonal orientation (30-60°). Both optoGEF and optoGAP embryos showed increased numbers of edges that form in an orientation similar to the initial orientation of AP edges (60-90°). n=3-5 embryos per genotype, 10-25 junctions/embryo. **(F-H)** Still images from time-lapse movies showing the progression of a cell rearrangement over time. optoGEF embryos showed increased numbers of newly formed edges oriented diagonally (not parallel) to the AP axis compared to controls. optoGAP embryos showed increased times to resolve a vertex during cell rearrangement compared to control or optoGEF embryos. Bars, 5 μm.

We also analyzed how the altered Rho1 activity in photoactivated optoGEF and optoGAP embryos influenced the spatial organization of cell rearrangements. In control embryos, 97% of vertices formed through contraction of an AP cell edge went on to resolve and form a DV cell edge, to produce a productive rearrangement (Fig. 6*D,F*) consistent with previous studies of cell rearrangement (Bertet et al., 2004; Blankenship et al., 2006; Kasza et al., 2019, 2014; Paré et al., 2014). The remaining 3% of vertices failed to resolve within 10 min after the formation of the vertex. In contrast, the percentage of vertices that resolved to form DV edges was reduced to 82% in photoactivated optoGEF embryos (*P* = 0.14) and 60% in photoactivated optoGAP embryos (*P* = 0.005) (Fig. 6*D*). In optoGEF embryos, the remaining 18% of vertices resolved to reform AP cell edges, resulting in unproductive cell rearrangement. In optoGAP embryos, 16% of vertices resolved to form AP cell edges and 24% did not resolve. In addition, we observed that new junctions formed during cell rearrangement in optoGEF embryos exhibited a disruption in new edge orientation. We observed a higher percentage of new edges forming at 30-60° (“diagonal” to the AP axis) compared to in control and optoGAP embryos (Fig. 6*E,G*), suggesting that the spatial orientation of rearrangements was disrupted by these perturbations.

Thus, inactivation of Rho1 in optoGAP embryos resulted in slower and significantly fewer completed cell rearrangements compared to control embryos, consistent with both a reduction in the overall levels and a disruption to the anisotropy of the myosin-generated forces driving tissue flow. In contrast, activation of Rho1 in optoGEF embryos resulted in less severe defects in the number and speed of rearrangements, but the rearrangements that did initiate were poorly oriented, which may explain why rearrangements in these embryos were less productive at producing tissue-level elongation (c.f. Fig. 1*C,F*). The increased cell rearrangement rates in optoGEF compared to optoGAP perturbations, might reflect the enhanced pulsatile medial-apical myosin contractility in these cells, which have the potential to promote cell rearrangements, especially in the absence of strong myosin planar polarity patterns. Consistent with this picture, cell rearrangements in optoGEF cells with enhanced medial myosin, which produced more isotropic mechanical tensions in the tissue, were in effect more “randomly” oriented within the tissue.

Taken together, these finding demonstrate that optogenetic activation or inactivation of Rho1 across the apical surface of the germband tissue produces distinct defects in the cell behaviors that contribute to convergent extension, directly linking patterns of upstream Rho1 activity and downstream actomyosin contractility to the ability of the tissue to deform and flow through cell rearrangements.

## DISCUSSION

Rapid changes in actomyosin localization and dynamics are coordinated with morphogenetic events in developing embryos, but it is not understood how actomyosin activity influences both tissue mechanical properties and driving forces in a manner that promotes rapid cell rearrangements and tissue remodeling. Here we used optogenetic tools to manipulate the patterns of actomyosin activity at the apical surface of the *Drosophila* germband epithelium during the converging and extending tissue flows of body axis elongation. We found that optogenetic activation or deactivation of Rho/Rho-kinase signaling both disrupted oriented cell rearrangements and tissue flow, leading to reduced tissue fluidity despite distinct effects on myosin localization and activity patterns. Based on laser ablation analyses to probe forces and cell packing analyses to infer tissue mechanical properties, our results indicate that actomyosin activity influences both the anisotropic forces driving flow and the mechanical properties of the tissue that characterize resistance to flow. These dual roles for actomyosin in tissue flow mechanics highlight the complex nature of the mechanisms that regulate actomyosin activity to control the cell behaviors that produce tissue shape and structure.

Cell rearrangements within a tissue can lead to macroscopic tissue-level shape changes if rearrangements are spatially and temporally coordinated across the tissue, or alternatively, no tissue-level shape changes if rearrangements occur in a more randomly oriented fashion (Tetley and Mao, 2018). In wild-type embryos, oriented cell rearrangements are a major contributor to germband extension (Bertet et al., 2004; Blankenship et al., 2006; Irvine and Wieschaus, 1994; Zallen and Wieschaus, 2004). We found that optogenetically increasing (optoGEF) or decreasing (optoGAP) Rho/Rho-kinase activity across the apical surface of the germband led to similar decreases in tissue-level flow, even though more rearrangements were initiated in the optoGEF compared to optoGAP embryos. The presence of cell rearrangements in optoGEF embryos that do not productively drive tissue elongation indicates that uniform apical activation of Rho/Rho-kinase signaling altered the spatiotemporal organization of cell rearrangements. Our studies reveal that the perturbations in optoGEF embryos led to a rapid redistribution of myosin from a planar polarized junctional pattern to a more radial, medial-apical pattern, associated with cell rearrangements that were more likely to resolve in the wrong direction, consistent with prior findings that tension distributions affect the orientation of vertex resolution and new edge formation (Curran et al., 2017; Yu and Fernandez-Gonzalez, 2016). The poorly oriented rearrangements are distinct from the less productive cell rearrangements observed in the germband of embryos expressing phosphomimetic myosin regulatory light chains in which there were additionally cell edge contraction errors associated with aberrant junctional accumulation and medial depletion of myosin II (Kasza et al., 2014). By comparison, in the *Drosophila* pupal notum, a tissue that exhibits cell rearrangements but no macroscopic tissue-level deformation, myosin II is not planar polarized and cell rearrangements may be transient or occur in random orientations (Curran et al., 2017). Our findings reveal that planar polarized patterns of junctional myosin are required for rapid cell rearrangements *and* tissue-level elongation, whereas isotropic patterns of medial-apical myosin are sufficient to promote rearrangements but do not orient rearrangements to produce tissue-level flows in this context. Taken together, these findings highlight the key role of planar polarized junctional myosin activity in spatially organizing cell rearrangements to harness tissue fluidity for rapid tissue shape change.

Cellular actomyosin contractility contributes to tissue-level patterns of mechanical tension that can drive tissue flow and have the potential to additionally tune tissue mechanical properties. The decreased myosin planar polarity observed in photoactivated optoGEF and optoGAP embryos would be predicted to result in reduced anisotropy in forces along AP compared to DV cell junctions. Consistent with this, we found decreased anisotropy in tensions, supporting a strong link between internal myosin patterns and tissue tensions. However, we did observe residual mechanical anisotropy in some optoGEF and optoGAP embryos, which might result from differences in contractile activities of myosin localized at AP and DV junctions, external stresses, or geometric constraints. Reductions in tension anisotropy have also been observed in *bicoid nanos torso-like* mutant embryos lacking AP patterning signals (Fernandez-Gonzalez and Zallen, 2011), *torso* mutant + *eve* RNAi embryos in which internal forces associated with cell intercalation are disrupted and external forces associated with posterior midgut invagination are blocked (Collinet et al., 2015), and embryos expressing phosphomimetic myosin regulatory light chain variants with disrupted myosin planar polarity (Kasza et al., 2014). In particular, the tissue retraction patterns following ablation in optoGEF embryos resembled those in *bicoid nanos torso-like* mutant embryos, which also display increased myosin at the medial apical cortex (Fernandez-Gonzalez and Zallen, 2011). The ring-like medial myosin organization that we observed in optoGEF embryos is also reminiscent of the pattern in apically constricting cells under isotropic tension in the invaginating posterior midgut (Chanet et al., 2017). Our results indicate that the myosin accumulating at the apical cortex of germband cells in optoGEF embryos leads to relatively isotropic tissue behavior in terms of resistance to contractile tension. The disruption of anisotropic contractile tensions associated with the loss of junctional myosin planar polarity in optoGAP and optoGEF embryos suggests that the rapid and dramatic transformation of myosin-dependent forces contributes to the defects in tissue flow.

During early *Drosophila* development, the mechanical properties of the tissue are thought to be largely determined by the cells themselves. Although techniques to measure tissue mechanical properties have been developed (Bambardekar et al., 2015; Campàs, 2016; D’Angelo et al., 2019; Doubrovinski et al., 2017; Herrera-Perez and Kasza, 2018; Mongera et al., 2018; Petridou and Heisenberg, 2019; Serwane et al., 2016), direct characterization of epithelial tissue sheets in the developing *Drosophila* embryo remains a significant challenge. To gain insight into tissue mechanical behavior, theoretical models of epithelia have been used extensively (Alt et al., 2017; Atia et al., 2018; Bi et al., 2016, 2015; Farhadifar et al., 2007; Fletcher et al., 2014; Krajnc et al., 2018; Nagai and Honda, 2001; Rauzi et al., 2008; Staple et al., 2010; Wang et al., 2020; Yan and Bi, 2019). A growing body of research indicates that cell shapes and packings within epithelia are intimately linked with tissue mechanics (Bi et al., 2016, 2015, 2014; Duclut et al., 2021; Farhadifar et al., 2007; Jain et al., 2020; Mitchel et al., 2020; Park et al., 2015; Popović et al., 2021; Staple et al., 2010; Wang et al., 2020). Indeed, recent work from our group and others indicate that two metrics for cell packings, the cell shape index and the cell shape alignment index, can be used to predict the onset of rapid cell rearrangements within the germband and that this corresponds to the solid-to-fluid transition in the vertex model (Wang et al., 2020). Here, we found that either activation (optoGEF) or inactivation (optoGAP) of Rho/Rho-kinase signaling in the germband alters cell shapes and packings in the germband. The cell packings in the perturbed tissues suggest less fluid-like (more solid-like) tissue mechanical properties compared to wild-type embryos, consistent with the reductions in cell rearrangements observed in these perturbations. In *Xenopus* explants, actomyosin contractility has been shown to promote stiffening of the tissue (Zhou et al., 2009), while during zebrafish posterior body axis elongation, inhibition of myosin II activity is associated with a rigidification of the tissue (Mongera et al., 2018). In vertex models of epithelia, the preferred shapes of cells and the associated collective energy barrier to cell rearrangement have been proposed to depend on a balance between myosin-dependent contractile tension and E-cadherin-dependent cell-cell adhesion at cell contacts, with enhanced contractile tension predicted to increase rearrangement energy barriers and produce more solid-like tissues (Bi et al., 2015; Park et al., 2015). Our findings suggest that actomyosin activity influences germband tissue mechanical properties but that there is not a simple monotonic relationship between overall levels of myosin activity and tissue mechanical properties. Instead, our results indicate that the overall levels, junctional versus medial accumulation, and planar polarized patterns of myosin activity are all likely to influence the energy barriers to cell rearrangement that determine tissue mechanical behavior.

Recent theoretical and experimental studies have highlighted that the mechanical behavior and flow of tissues can depend on active fluctuations within cells, which promote cell rearrangements in tissues with finite energy barriers to cell rearrangements. Although we observed similar disruptions to myosin planar polarity and tension anisotropy in both the optoGEF and optoGAP perturbations, these two cases are distinguished by the different rates of cell rearrangements in the germband, with roughly 2-fold more rearrangements per cell per minute in optoGEF compared to optoGAP embryos. This is despite the fact that the germband in the optoGEF embryos is predicted to be *less* fluid-like and have *higher* energy barriers to rearrangement within the vertex model framework. A notable feature of these perturbations that appears to correlate with the observed rearrangement rates is the enhanced medial-apical myosin in optoGEF and decreased medial-apical myosin in optoGAP embryos. In prior work, we showed that the enhanced medial-apical myosin pool in the photoactivated optoGEF cells displays dynamic pulsing behavior that is correlated with apical cell area fluctuations (Herrera-Perez et al., 2021). Here, by studying retraction of the tissue in response to laser ablation of the medial myosin domain, we demonstrated that this enhanced medial-apical myosin pool produces enhanced active tensions. The presence of cell rearrangements in optoGEF embryos suggests that pulsatile contractile forces originating from the medial-apical domain of cells might suffice to promote cell rearrangements in the absence of planar polarized driving forces, though such rearrangements are more likely to resolve in the wrong direction. In wild-type embryos, previous studies have demonstrated roles for pulsatile myosin activity during cell edge contraction (Rauzi et al., 2010) and for medial myosin pulsation in the adjacent cells during new edge formation (Collinet et al., 2015; Yu and Fernandez-Gonzalez, 2016) to promote oriented cell rearrangement. Our results support the notion that pulsatile medial-apical contractility provides active tension fluctuations that can promote cell rearrangements and fluidize tissues, but that this is not sufficient to drive tissue-scale flow in the absence of anisotropic forces to spatially direct these rearrangements.

Even though optogenetic perturbations significantly altered actomyosin patterns in the germband, influencing cell packings, and producing tissues predicted to have increased energy barriers to cell rearrangement and less fluid-like properties, these perturbations were not sufficient to produce cell packings during germband extension characteristic of solid-like tissues. This raises the question of what additional molecular and mechanical factors are involved in controlling the solid-fluid properties of tissues. Future work will be needed to further explore how junctional and medial myosin pools work together to tune tissue mechanics in the germband and in other developing epithelia. In addition, the contributions of cellular adhesive machineries and their interplay with the actomyosin cytoskeleton will need to be investigated. Ultimately, to understand how emergent tissue mechanics and flows are regulated in space and time across the developing embryo to generate functional three-dimensional embryonic structure, we will need to identify the master upstream regulators that tune and coordinate force generation and tissue mechanical properties to achieve both the local cell structure and global tissue shape required for proper function. The approaches and findings we report here will be useful in this endeavor.

## MATERIALS AND METHODS

### Fly lines and genetics

Embryos were maintained at 23°C and experiments were performed at room temperature (~ 21°C). The fly stock w[*]; P[w+, UASp>CIBN-caax] was generated in this study by PCR amplification of the CIBN-caax fragment from plasmid pPW_CIBNcaax (gift from Stefano De Renzis). The fragment was cloned into the pENTR/D-Topo vector (Life Technologies) and recombined into UASp-attB destination vector (gift from F. Wirtz-Peitz) using the Gateway cloning system (Life Technologies) for subsequent expression using the Gal4/UAS system (Brand and Perrimon, 1993). The plasmid was inserted into the attP40 site on chromosome II to generate transgenic flies (Bestgene).

The fly stocks w*;; w+, UASp>mCherry-CRY2PHR-RhoGEF2 and w*;; w+, UASp>mCherry-CRY2PHR-RhoGAP71E were crossed with w*; P[w+, UASp>CIBN-pmGFP]/CyO; Sb/TM3,Ser (gift of Stefano De Renzis, EMBL, Heidelberg, Germany)(Guglielmi et al., 2015). The stocks w*;; w+, UASp> CRY2PHR-RhoGEF2 and w*;; w+, UASp> CRY2PHR-RhoGAP71E were crossed with w*; P[w+, UASp>CIBN-pmGFP]/CyO; Sb/TM3,Ser (gift of Stefano De Renzis, EMBL, Heidelberg, Germany) or w*; w+, UASp>CIBN-caax;. The constructs were expressed using the maternal α-tubulin matα-tub15 or matα-tub67 Gal4-VP16 drivers (mat67, mat15, gift of D. St Johnston).

To visualize myosin II, mat15 and sqh>sqh-mCherry (BDSC 59024, donated by Beth Stronach and Elane Fishilevich) were recombined on chromosome III. For Rho-kinase (Rok) visualization, mat15 and sqh>GFP-Rok^K116A^ (gift of Jennifer Zallen) (Simões et al., 2014) were recombined on chromosome III. To protect the embryos from light, crosses were maintained in the dark, and fly sorting was performed in the dark on a stereoscope equipped with Red 25 Kodak Wratten Filter.

Embryos were the progeny of females of the following genotypes:

UASp>CIBN-pmGFP/mat67 (II); sqh>sqh-mCherry/+ (III)
sqh> GFP-Rok^K116A^ (III)
UASp>CIBN-pmGFP/+ (II); UASp>mCherry-CRY2PHR-RhoGEF2/mat15 (III)
UASp>CIBN-pmGFP/+ (II); UASp>CRY2PHR-RhoGEF2/sqh>sqh-mCherry, mat15 (III)
UASp>CIBN-caax/+ (II); UASp>CRY2PHR-RhoGEF2/ sqh>GFP-Rok^K116A^, mat15 (III)
UASp>CIBN-pmGFP/mat67 (II); UASp>mCherry-CRY2PHR-RhoGAP71E/mat15 (III)
UASp>CIBN-pmGFP/mat67 (II); UASp>CRY2PHR-RhoGAP71E/sqh>sqh-mCherry, mat15 (III)
UASp>CIBN-caax/mat67 (II); UASp>CRY2PHR-RhoGAP71E/ sqh>GFP-Rok^K116A^, mat15 (III)

Fly stocks used in this study:

w*; w+, UASp>CIBN-pmGFP/CyO; Sb/TM3,Ser (Stefano De Renzis, EMBL, Heidelberg, Germany)
w*; w+, UASp> CIBN-caax; (this study)
w*;; w+, UASp>mCherry-CRY2PHR-RhoGEF2/TM3,Sb
w*;; w+, UASp>CRY2PHR-RhoGEF2/TM3,Sb
w*;; w+, UASp>mCherry-CRY2PHR-RhoGAP71E/TM3,Sb
w*;; w+, UASp>CRY2PHR-RhoGAP71E/TM3,Sb
w*;; sqh>sqh-mCherry
w*; w+, UASp>CIBN-pmGFP, mat67; sqh>sqh-mCherry
w*;; mat15
w*; mat67; mat15
w*;; sqh>sqh-mCherry, mat15
w*; mat67; sqh>sqh-mCherry, mat15
w*;; sqh>GFP-Rok^K116A^
w*;; sqh>GFP-Rok^K116A^, mat15
w*; mat67; sqh>GFP-Rok^K116A^ (III)

### Embryo preparation

Embryos were prepared and collected in a dark room illuminated with red light. Embryos in early stage 6 were dechorionated with 50% bleach for 2 min, washed with water, and mounted using a mixture 50:50 of halocarbon oil 27:700 on a custom-made imaging chamber between an oxygen-permeable membrane (YSI Incorporated) and a glass coverslip.

### Imaging

Embryos were positioned ventrolaterally for observation of the germband. Photoactivation and imaging was performed with a Zeiss LSM 880 confocal microscope with Airyscan detector and equipped with an Argon laser for 488 nm excitation and a diode laser for 561 nm excitation (Carl Zeiss, Germany). Imaging was performed with C-Apo 40X/1.2 NA water-immersion or Plan-Apo 63X/1.40 NA Oil-immersion objectives (Carl Zeiss, Germany). The bright field illumination light was filtered through a Red 25 Kodak Wratten Filter to prevent unwanted photoactivation. Image acquisition was performed with ZEN black software (Carl Zeiss, Germany) using the standard LSM mode unless otherwise noted. For myosin II and Rok imaging, the Airyscan FAST module was utilized. Images were acquired as 10 μm z-stacks with a 1 μm z-step (40X objective, LSM mode) or a 0.7 μm z-step (63X objective, Airyscan FAST), beginning at the apical side of the tissue. Z-stacks were acquired every 15 s. Photoactivation was achieved by scanning with blue laser light (λ = 488 nm) over the same ventrolateral region of the germband that was being imaged, at every imaging z-plane and time-step.

### Image analysis

Confocal time-lapse movies of the cell outlines visualized using CIBN-pmGFP, were projected using the ImageJ distribution Fiji software (Rueden et al., 2017; Schindelin et al., 2012) and used to calculate tissue elongation during body axis elongation using the particle image velocimetry (PIV) software PIVlab version 1.41 (Thielicke and Stamhuis, 2014) in MATLAB as previously described (Kasza et al., 2014; Thielicke and Stamhuis, 2014). To quantify vertex coordination number and number of rearrangements per cell, movies with fluorescently-tagged cell membranes were segmented using the SEGGA tissue segmentation and analysis software (Farrell et al., 2017). The instantaneous rate of rearrangements was calculated as a moving average of the cell rearrangement rate over 1.5 min windows.

### Signal intensity quantification

For myosin II analysis, still frames shown and confocal movies analyzed correspond to the maximum intensity projections of 5 μm at the apical side of the tissue, starting apically. Images were processed and analyzed using the ImageJ distribution Fiji software (Rueden et al., 2017; Schindelin et al., 2012). The mean junctional myosin intensity at each cell edge was quantified manually using 6 pixel-wide lines (excluding vertex regions) in a 40 μm square region, and the intensity and orientation of each edge was obtained. Medial apical myosin intensity was quantified manually as the mean intensity in a polygon manually drawn just inside the cell membrane outline. At least 10 cells and 40 junctions were analyzed per embryo (Fig. 3). Planar polarity was obtained as the ratio of the mean intensity of AP (“vertical”) edges to the mean intensity of DV (“horizontal”) edges for each embryo (Fig. 3).

### Cell rearrangement analysis

To quantify the number of rearrangements per cell (Fig. 1), movies with fluorescently-tagged cell membranes were segmented using the SEGGA tissue segmentation and analysis software (Farrell et al., 2017). To analyze the speed and orientation of rearrangements (Fig. 6), cell rearrangements were manually analyzed for 10-25 vertical cell edges per embryo (n=48-69 edges in total from 3-5 embryos per genotype). AP edges were selected at the beginning of axis elongation, and their fate was tracked over 30 min. Edge contraction rate was calculated from changes in edge length during the 2 min before vertex formation. New edge growth rate and orientation were calculated over the 2 min following vertex resolution. Vertex lifetime corresponds to the time between when the edge contracts to a vertex and when the new edge is > 1 μm long. Vertices that did not resolve corresponded to AP edges that contracted to a vertex but did not resolve to form a new edge within the tracking time-window (vertices that formed >20min after the start of the analysis window were not included in the analysis). Vertices that resolved in the wrong direction correspond to AP edges that contracted to a vertex and then resolved to reform an AP cell edge instead of forming a new contact between dorsal and ventral cell neighbors.

### Laser ablation

Samples for ablation studies were prepared in a dark room illuminated with red light. Embryos were dechorionated and mounted in a 50:50 mixture of halocarbon oils 27 and 700 within a custom-made imaging chamber. Ablations were performed using a Chameleon Ultra2 laser set to 710 nm on a Zeiss LSM 880 confocal microscope equipped with a Plan-Apo 63X/1.4 NA oil DIC M27 immersion objective. Embryos expressing optogenetic tools were photoactivated with blue light from an Argon laser (λ = 488 nm) at every z-plane (10 μm z-stacks with 1 μm steps) every 15 s per stack during the 5 min prior to ablations. The following laser settings were used for ablation: 60% laser power, 1 iteration, scan speed of 0.0143 ms per pixel, 1.3 μm line. After ablation, single z-plane images were acquired every 1 s for at least 30 s. For vertex retraction measurements following ablation of junctions, the distance between the vertices connected to the ablated edge were manually measured over time. To generate tissue displacement maps, displacements were quantified by particle image velocimetry (PIV) analysis (Matlab PIVlab version 1.41) (Thielicke and Stamhuis, 2014) using a 5 by 5 grid (3.3 μm between grid points) in a 174 μm^2^ region surrounding the ablation point over the course of 30 s following ablation. For radial plots, mean displacements were calculated in 60° angular bins along the anterior, posterior, dorsal, or ventral tissue axes, taking the ablation point as the origin.

### Cell shape index and cell shape alignment analysis

Cell outlines were segmented using the SEGGA tissue segmentation and analysis software (Farrell et al., 2017). For each segmented cell within a specified region of interest, the perimeter (*P*) and area (*A*) was measured and the ratio 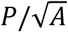 was calculated. The average shape index 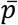 is the average value of 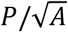 for all cells. To account for effects of cell packing disorder, the corrected shape index 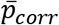 was calculated by subtracting the term (*z*-3)/*B* from 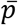, where *z* is the vertex coordination number (average number of cells meeting at a single vertex) and *B* is 3.85 (Yan and Bi, 2019), as previously described (Wang et al., 2020). To calculate the cell shape alignment *Q*, the tissue was triangulated as previously described (Merkel et al., 2019, 2017; Wang et al., 2020). For each triangle, an alignment tensor is calculated, and *Q* is the magnitude of the area-weighted average of the alignment tensors as described in (Merkel et al., 2019, 2017; Wang et al., 2020). As previously described, the predicted solid-fluid transition line is given by 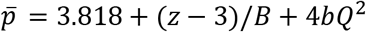, or alternatively, 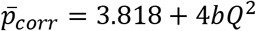 (Wang et al., 2020), where *b*=0.6 ± 0.2 based on previously reported vertex model simulation results (Merkel et al., 2019).

### Statistical analysis

Data are presented as mean values +/− the standard error of the mean (SEM). The nonparametric Kruskal-Wallis test with post-hoc Dunn test was used to determine P-values unless otherwise noted.

## ACKNOWLEDGEMENTS

We thank Nandan Nerurkar for use of and assistance with microscope and laser facilities for ablation studies, Stefano De Renzis for the UASp>CIBN::pmGFP fly stock, Jennifer Zallen and Frederik Wirtz-Peitz for the UASp-attB destination vectors, the Bloomington Drosophila Stock Center (BDSC) for fly stocks, and members of the Kasza Lab for helpful discussion and comments on the manuscript. This work was supported by the National Institute of General Medical Sciences of the National Institutes of Health Award Number R35GM138380 (to K.E.K.); Eunice Kennedy Shriver National Institute of Child Health & Development of the National Institutes of Health Grant F31HD105405 (to C.C); National Science Foundation Graduate Research Fellowship (to A.B.D.). M.H.P holds a Career Award at the Scientific Interface from the Burroughs Wellcome Fund. K.E.K holds a Career Award at the Scientific Interface from the Burroughs Wellcome Fund, a Clare Boothe Luce Professorship, and a Packard Fellowship.

